# Towards faster identification of MRSA and MSSA: analysis of calorimetric curve parameters from large hospital bacterial collections

**DOI:** 10.1101/2023.11.28.568793

**Authors:** Sara Zunk-Parras, Andrej Trampuz, Flor I. Arias-Sánchez

## Abstract

There is a need to develop faster yet precise diagnostic tools for the identification of infectious agents and their levels of antimicrobial susceptibility. One such tool is calorimetry, as previous studies show that bacteria display unique signatures of calorimetric/metabolic activity that such signatures can be used for pathogen identification under controlled laboratory conditions. However, the level of variation of these unique signatures remains largely unexplored. Here, we use real-time isothermal microcalorimetry to disentangle the inter- and intra-species metabolic differences between *S. aureus* and *S. epidermidis*. We worked with a large collection of bacterial isolates obtained from patients with prosthetic joint infections as well as reference laboratory strains. We analyzed their calorimetric profiles and decomposed the curve parameters derived from them. As a result, we identified six calorimetric parameters that are useful for identification at the intra-species level, with a particular focus on MRSA. Furthermore, we found important differences between the thermograms of ATCC laboratory strains when compared against model thermograms generated from all our clinical isolates. These results indicate that accounting for metabolic variability can impact diagnosis precision. Finally, we discuss our findings and suggest ways to optimize calorimetric diagnostics and treatment approaches.

## INTRODUCTION

Antimicrobial resistance (AMR) is a threat to global health given the estimation that it will cause more than 10 million deaths by the year 2050 (1, 2). New technologies that reduce diagnostic times and allow for simultaneous antimicrobial susceptibility testing in both planktonic and biofilm forms, are crucial for the current wide-ranging health (3). Current routine diagnostics such as microbiological cultures in nutrient media and polymerase chain reactions (PCRs), need between 24 h (4) and up to five days (5), depending on the test and the biological sample to be tested. Faster molecular-based diagnostic techniques are arising: such as direct MALDI-TOF-MS (6, 7) or isothermal microcalorimetry which detects the growth-related heat release of metabolically active cells in both planktonic and biofilm forms (5). The detection limit of isothermal microcalorimetry is 200 nW, meaning that only as few as 100,000 bacteria are needed for detection, since a typical growing bacterium generates around 2 pW (8). This makes this real-time detection technique highly sensitive and useful in multi-variable studies, which consequently is an adequate tool for susceptibility testing of various antimicrobials (4, 9, 10).

Isothermal microcalorimetry has shown to be useful in the diagnosis of synovial fluid infections (5, 11), as well as in the detection of infected tissues (4) with similar detection levels as bacterial cultures but with the advantage of giving signals of bacterial growth in a shorter period of time. However, the fact that it measures a nonspecific signal based on the heat flow occurring inside the calorimetric vials, creates a potential limitation for pathogen species identification (4, 10). Therefore, further improvements of this technique need to be implemented in order to exploit its possible use as a diagnostic tool.

Recent studies show that laboratory strains of various bacterial species generate different and unique thermogram profiles (4, 5). Baldoni et *al* showed the differences in the heat flow curves generated by MRSA and MSSA strains (12). Also, Grütter et *al* investigated detection and drug susceptibility testing in two different strains of *Neisseria gonorrhoeae* through isothermal microcalorimetry, and highlighted the differences found in the thermograms between strains of the same organism (13). These results suggest that characterization of the thermogram profiles could facilitate the identification of the microorganism causing an infection, including its level of susceptibility to antibiotic treatment.

However, the limitation of these studies is that they focus on few bacterial strains. They often focus on ATCC laboratory strains which are adapted to the laboratory conditions, therefore ignoring the species diversity that is usually found in nature and the hospital environment (14-18). Studies addressing the levels of inter and intra-species diversity and their impact on infection identification using calorimetry are still lacking. Addressing this gap in knowledge is important to better understand whether thermograms can reliably be used as a diagnostic tool. Here, we aimed to evaluate and quantify the metabolic differences at the inter- and intra-species level and identify key curve parameters that could allow for quick diagnosis. We focus on *Staphylococcus aureus* and *Staphylococcus epidermidis*, due to their prevalence in the clinic (19). We study a large collection of 171 clinical isolates and 9 ATCC laboratory strains and use analysis of thermogram data to (i) address the level at which inter- and intra-species diversity affects pathogen identification; (ii) identify the calorimetric parameters useful for bacterial identification and distinction of resistance patterns; and finally (iii) we propose a workflow to use clinical strain collections for the production of model curves, which can hopefully aid in the improvement of calorimetry as a diagnostic tool.

## MATERIALS AND METHODS

### Strains and culture conditions

We evaluated two large collections of clinical isolates obtained from 171 partially immunocompromised patients treated for various types of infections at Charité University Hospital Berlin (Center for Musculoskeletal Surgery) between January 2015 and July 2019, including prosthetic joint infections, implant-associated infections and wound infections. The first BioBank collection corresponds to 84 clinical isolates obtained from infections caused by *S. aureus*, while the second collection corresponds to 87 clinical *S. epidermidis* clinical isolates. All strains were previously identified via MADI-TOF-MS, as part of the routine diagnostic procedure at Labor Berlin – Charité Vivantes GmbH.

For our study, we obtained pure cultures each of these isolates by streaking in agar media and isolating single colonies. After two rounds of single colony isolation, we grew overnight cultures from just one colony to ensure purity of our cultures. We produced pure stocks of each isolate, by mixing 700 μL of the overnight cultures with 700 μL of glycerol 50%. In each collection, we included ATCC strains as controls, three for *S. epidermidis* (*ATCC 20048, ATCC 3269, ATCC 18857*) and six for *S. aureus*, three of which were MRSA (*ATCC 43300, MRSA Col, MRSA Mu3*) and the other three were MSSA (*ATCC 29231, ATCC 104437, ATCC 105272*). The *S. aureus* strains were grown in Mannitol Salt Agar, (Oxoid, CM0085) while *S. epidermidis* in Tryptic Soy Agar (US Biological Life Sciences, L22051322; Millipore, 9002-18-0), both at 37 ºC.

### Determination of metabolic activity using isothermal microcalorimetry

We obtained the calorimetric profiles of both BioBank collections by isothermal microcalorimetry, using a calScreener device (Symcel AB) according to the manufacturer’s instructions (20). We grew each strain in two rich nutrient media suitable for the growth of our species (TSB and MHB, US Biological Life Sciences, L22051322; Carl Roth, X927.1). The initial inoculum from each strain was standardized by diluting pure overnight cultures to 0.5 McFarland, before a further 1:1000 dilution in the used medium. Each sample was loaded in separate vials with a total volume of 350 μl. We incubated at 37 °C and performed calorimetric measures every 10 minutes during 24 h. Thermograms were exported both as a matrix file with one data point every 10 min and as calorimetric curves.

### Analysis of inter and intra species diversity on diagnosis resolution

First, we evaluated which nutrient media permitted a better comparison of our bacterial groups, meaning which medium included more calorimetric parameters with significant differences when comparing the species grown with it. We tested these differences through Multivariate Analysis of Variance (MANOVA) (see supplementary material, **dTable S2**). Then, we used the data obtained from this specific medium to identify the calorimetric parameters that allowed species discrimination, taking the previous results from MALDI-TOF-MS analyses. For this, we analyzed the thermograms of each isolate through PCAs using R version 4.0.3 (21, 22). Separated clusters for each species in the PCAs would mean that the variance of the whole data set could be explained by the first two components generated, while overlapping clusters in the PCAs would represent that these components are not able to appear for the variance of our collection. In this last scenario, we would need to further analyze the thermograms of each isolate, in order to find calorimetric differences that would permit bacterial identification.

### Identification of curve parameters suitable for quick diagnosis

We decomposed the thermogram data into a corresponding set of 11 calorimetric parameters per curve (listed in **Table 1**) using the online tool calData (Symcel AB). Each of these parameters represents a specific quality of the curve (as shown in **Figure 1)**. To identify the parameters with the most predictive power for microbial pathogen identification, we performed an analysis of each parameter through independent tests. We tested each parameter independently, looking for significant differences at the inter-species level (comparing S. epidermidis against MSSA and MRSA respectively) and the intra-species level (comparing the pair MRSA against MSSA). For this, we used the non-parametric Kruskal Wallis test, followed by a Wilcoxson signed-rank test with a Bonferroni correction for multiple pairwise comparisons.

**Table 1.**
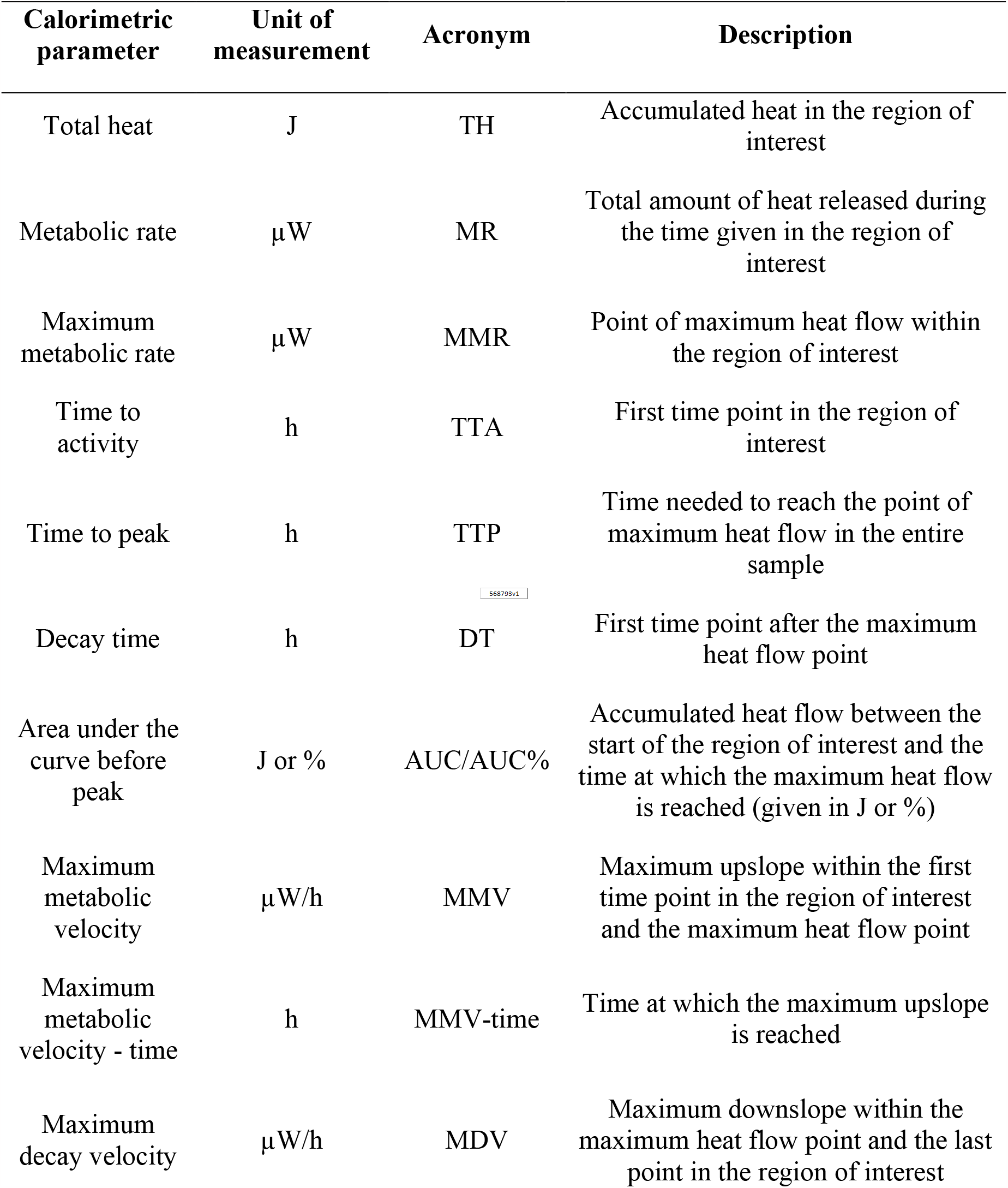
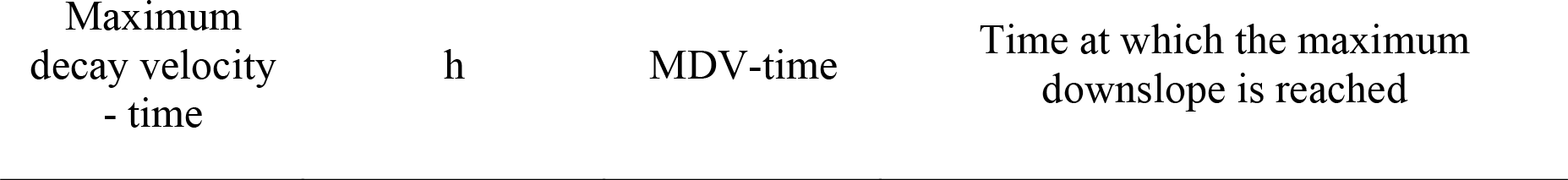
List of calorimetric parameters that can be exported from the calData tool. The description of the parameters has been taken from Symcel AB. For a visualization of the parameters, see **Figure 1**.

**Figure 1.**
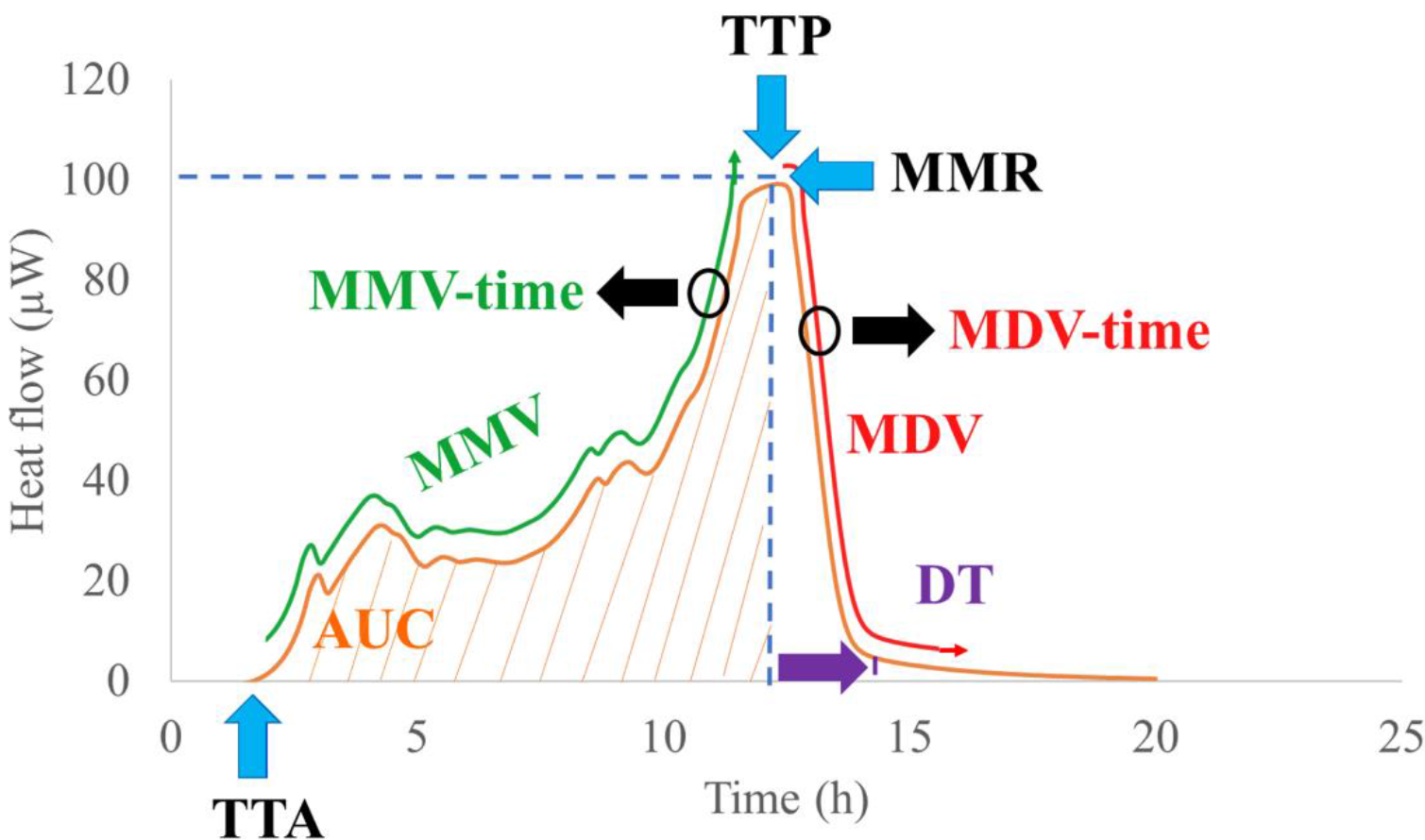
Illustration of the calorimetric parameters present in a thermogram. The maximum metabolic rate (MMR) is the highest heat point released, and the time to peak (TTP) is the time needed to reach this point. The time to activity (TTA) is the time needed to detect the start of the heat release. The area under the curve before the peak (AUC) is the total heat released from the starting point to the maximum metabolic rate. The decay time (DT) is the time needed from the maximum metabolic rate to the end of the decay signal. The maximum metabolic velocity (MMV) and maximum decay velocity (MDV) are the maximum acceleration during the rise and decay signals, respectively. The circles highlight the point where there is a maximum acceleration in the metabolic (MMV-time) and decay velocity (MDV-time), which are measures of time.

### Determination of reference thermograms: ATCC curves vs model curves

We tested the validity of using ATCC thermograms as standard references for pathogen identification, as opposed to using model curves constructed from all our data. First, we estimated the average calorimetric values per time point for each isolate when grown in TSB. We used these average values to produce model calorimetric curves for each species. We then compared these model curves against their respective ATCC curve. High similarity between the curves would be an indication that ATCC is a good standard reference for clinical diagnosis. On the contrary, low similarity would indicate that ATCC references are likely to mislead the identification of clinical samples. Additionally, we generated their corresponding calorimetric parameters, also obtained from the median values of all the calorimetric parameters obtained from our entire collection, which allowed a better comparison of both curves.

## RESULTS

### Accounting for strain diversity does impact calorimetric diagnostic resolution

PCA analysis of our 24h-thermograms (n=371) revealed a high level of overlap between clusters (**Figure 2a**). The two first component explained just 63% of the variance in our data (PC1 = 35.8%, PC2 = 27.6%; **Figure 2b**). To see the importance of each component, see **Table S1**. These results indicate a high level of variation in our clinical isolates, both at the inter- and intra-species level. Therefore, no separated clusters were differentiated due to media or species. Nevertheless, this did not interfere with the possibility to differentiate species and isolates through calorimetry.

**Figure 2.**
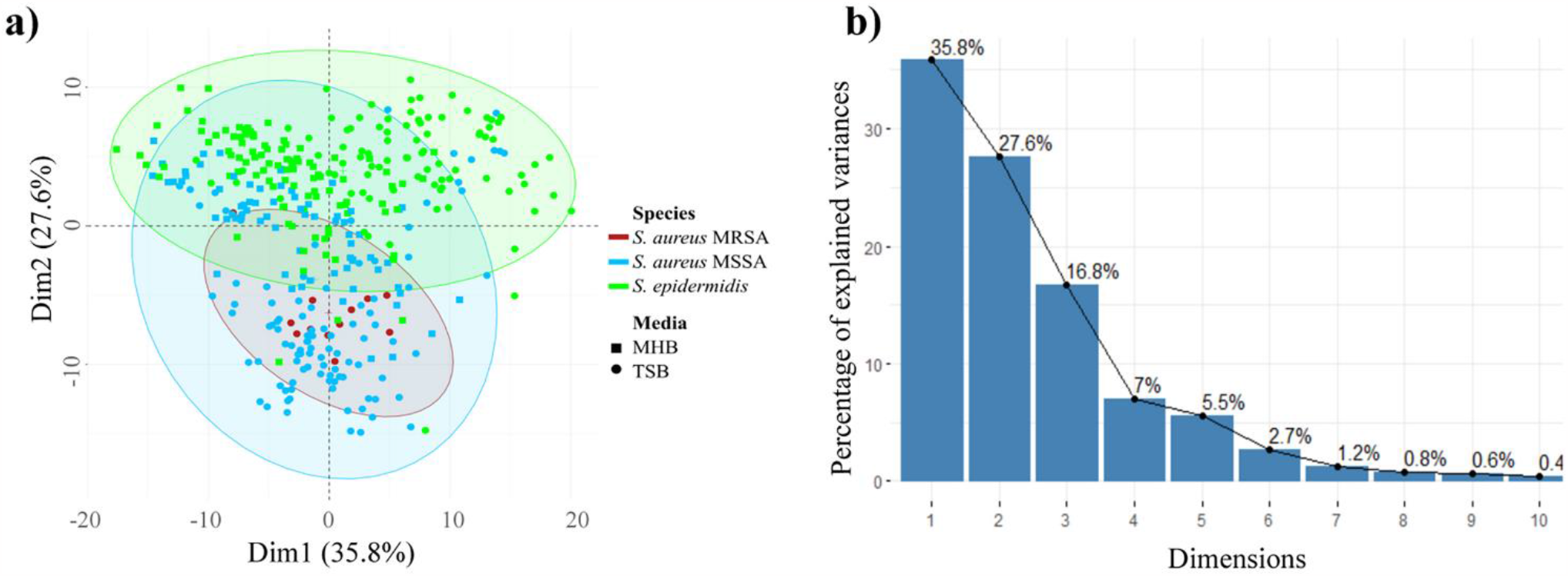
PCAs of thermogram data from clinical isolates of *S. aureus* and *S. epidermidis* grown in MHB and TSB media. (**a**) Biplot of the two first components generated with the thermograms of all the clinical isolates. Prediction ellipses are such that with probability 0.95, a new observation from the same group will fall inside the ellipse. N = 371 data points. (**b**) Scree-plot showing eigenvalues of each principal component. Figure and histogram generated with R version 4.0.3 (22).

### Identifying MRSA strains through parameter analyses

Analyses of each of our 11 calorimetric thermogram parameters in each bacterial collection (shown in **Table 1)** revealed that six of them show statistically significant differences between the three groups tested, making them suitable for determination of bacterial species and resistance patterns identification. These six parameters are time to peak, time to activity, maximum metabolic rate, maximum metabolic velocity, maximum metabolic velocity (time) and maximum decay velocity (**Figure 3**). The rest of the parameters are represented in **Figure S1**.

**Figure 3.**
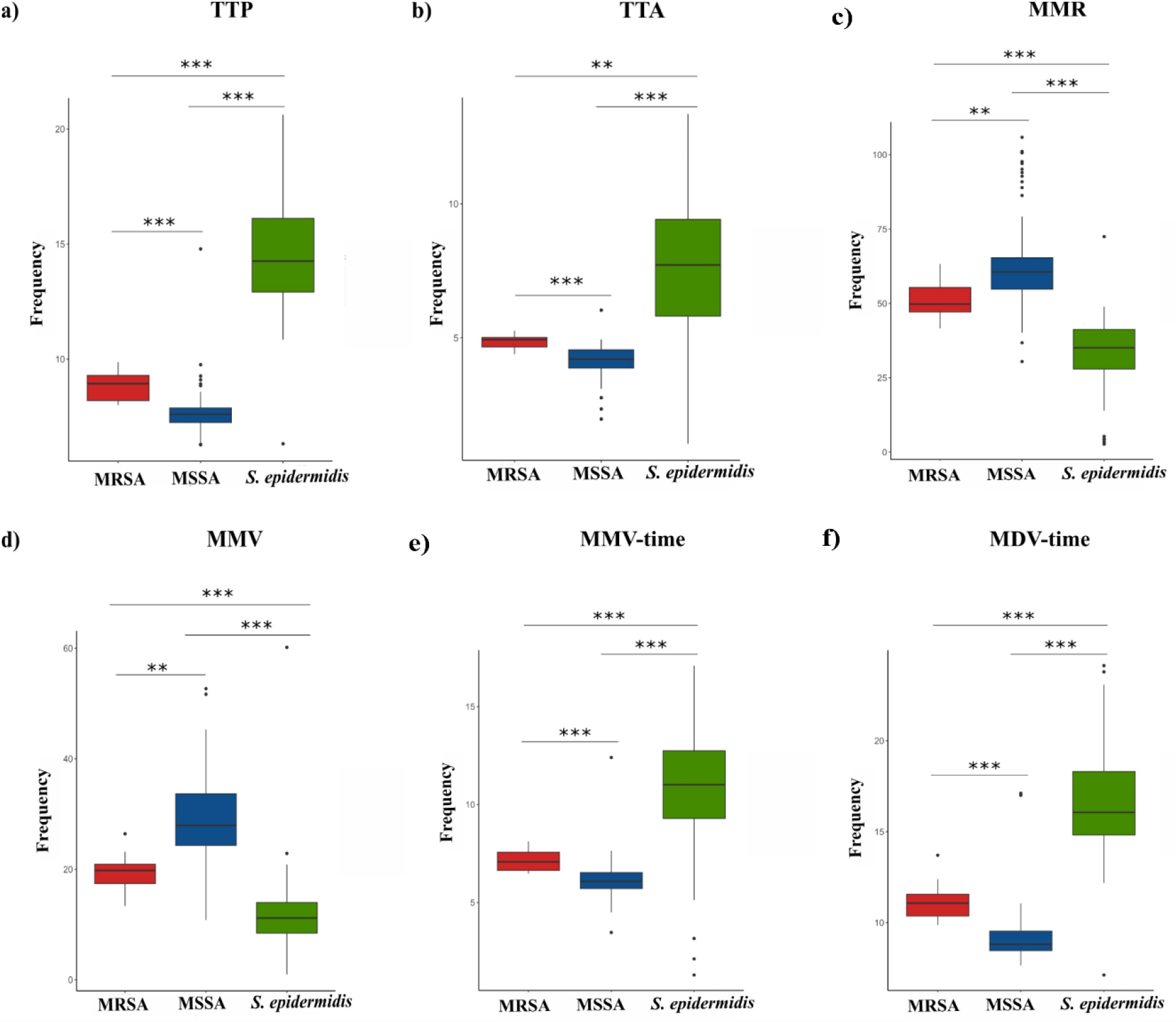
Calorimetric parameters that allow inter and intra species differentiation. Within each parameter, we perform pairwise comparisons between MRSA, MSSA and *S. epidermidis* isolates tested when grown in TSB. A total of 11 parameters were compared through Wilcoxson test with a Bonferroni adjustment to control for multiple comparisons. Significant differences between the parameters are represented by asterisks, where * represents p < 0.05, ** corresponds with a p < 0.01 and *** show a p < 0.001.

### Model curves may allow detection of MRSA infections in less than 8 h

#### Prediction curves that aid real-time diagnosis in the clinic showed differences between strain types

We found important differences in our comparison of model curves against ATCC curves, both at the inter- and intra-species level (**Figure 4a**). The model curves of *S. epidermidis* show higher discrepancies between ATCC strains and clinical isolates, as the curves present a different shape, leading to more remarkable divergence between the calorimetric parameters. On the other hand, the ATCC curves are very similar between them, especially regarding to the shape of the curve.

**Figure 4.**
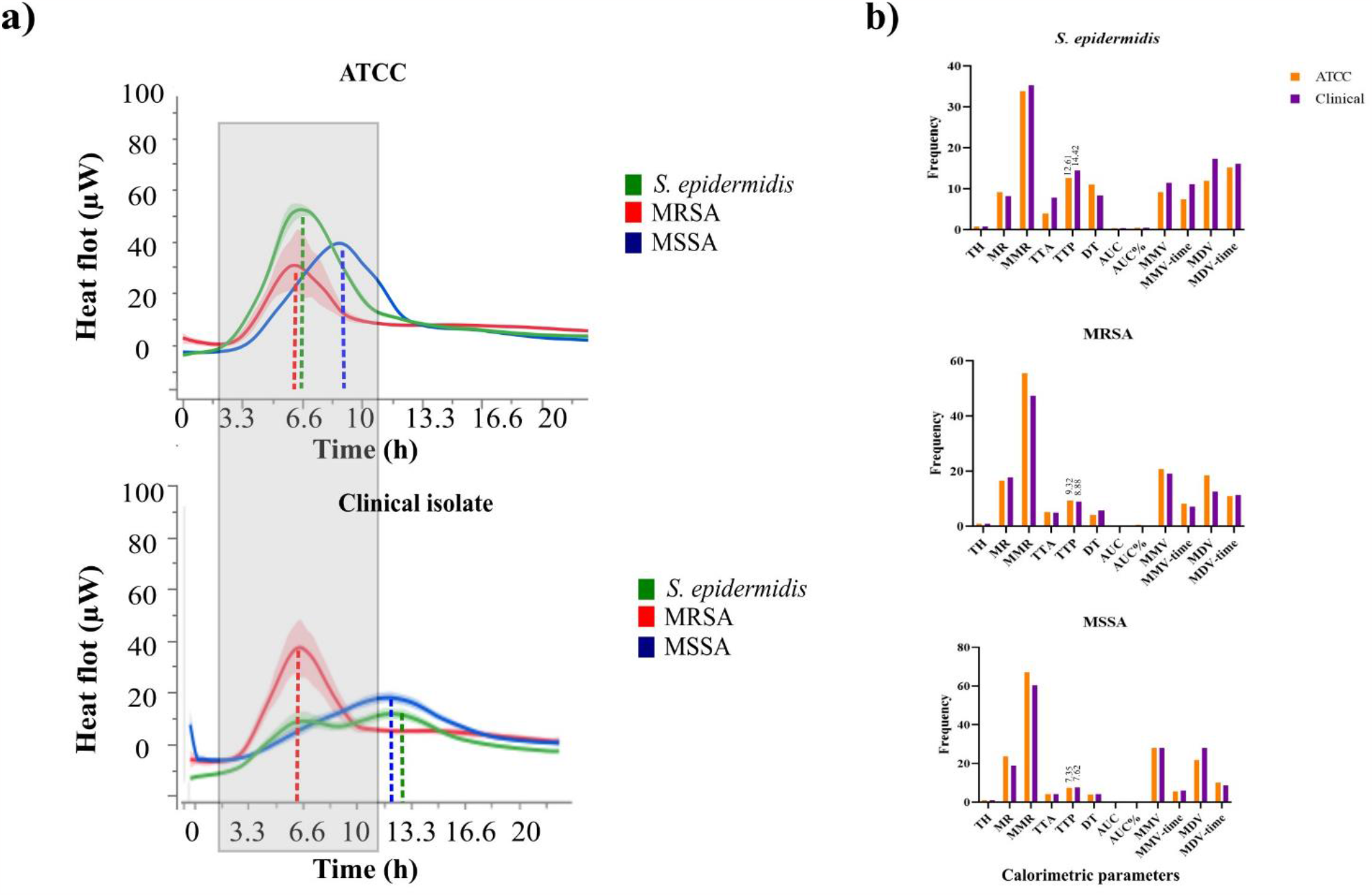
Metabolic differences between standard laboratory strains and clinical isolates. (**a**) Prediction curves built with the mean heat values exported for each strain. The time to peak is represented with dashed lines for each strain. Highlighted in grey is the time period where all the ATCC strains present their time to peak, while in the clinical isolates the same period includes exclusively the same parameter for MRSA isolates. (**b**) Differences in the mean values of the calorimetric parameters of each species. Mean values of the ATCC strains are shown in orange, while those from clinical isolates are shown in purple. The mean values of the time to peak for each strain are highlighted above the bars.

By comparing the calorimetric parameters of both strain types (**Figure 4b**) we found that the time to peak is delayed in *S. epidermidis* and MSSA isolates (14.42 h and 7.62 h, respectively) compared to the ATCC strains (12.61 h and 7.35 h, respectively), but strikingly is reduced in MRSA isolates, where the time to peak is at 8.88 h instead of at 9.32 h in the ATCC strains. These results show that the differences in the thermograms from clinical isolates of different species, as well as those from strains with different resistance profiles, can be identified by isothermal microcalorimetry but do not concord with those form the standard laboratory strains.

## DISCUSSION

We have focused on several aims in this study. First, we wanted to investigate the diversity of our bacterial collections to evaluate its impact on bacterial identification. Our PCAs derived from the thermograms of clinical isolates of *S. aureus* and *S. epidermidis* showed a high variability between these species and, therefore no clusters were differentiated. We also focused on the identification of calorimetric parameters that could be useful for pathogen detection and the distinction of their resistance patters. For this, we obtained the parameters of all our strains and performed pairwise comparisons. We identified six parameters that can be useful for bacterial identification. Furthermore, the same parameters showed statistically significant differences between the *S. aureus* isolates that are methicillin-resistant and methicillin-sensitive (**Figure 3**), showing the potential of this tool to also detect resistance patterns. Lastly, we wanted to propose a workflow to implement calorimetry as a diagnostic tool (**Figure 5**), for which we generated model calorimetric curves that are helpful for faster identification of bacterial growth and resistance patterns.

**Figure 5.**
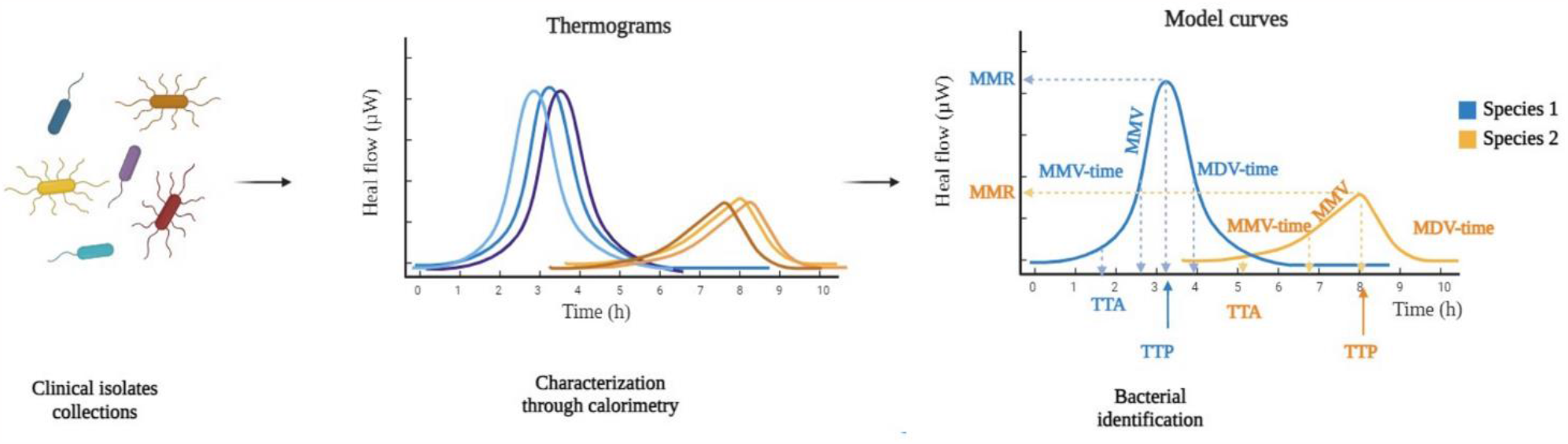
Proposed method of calorimetric analysis to produce model curves from clinical isolates’ collections. The characterization of these collections through isothermal microcalorimetry permits the generation of model calorimetric curves. The analysis of the derived calorimetric parameters can ultimately be useful for bacterial identification and, therefore, diagnosis of infectious diseases. Figure created with BioRender.com (2023).

The overlapping clusters that we obtained from our PCAs (**Figure 2**) highlight the importance of considering bacterial diversity in diagnostic studies. We conducted the study with 171 clinical isolates of two closely related species, hence the overlap in our biplots was anticipated. We would also expect that the more distant the species are, the less overlap we would observe in the clusters. To test the reproducibility of these results and extrapolate the use of microcalorimetry as a diagnostic tool, further research should be conducted with other staphylococci before expanding it to other bacterial species. We would need to examine the possibility of using isothermal microcalorimetry for the identification of other species, including gram-negative bacteria, as well as other microorganisms such as fungi. Moreover, we would need to also expand the study of resistance patterns, further than resistance to methicillin. This is something that remains to be explored in the future.

Given the strong overlap, we focused on the identification of specific parameters that could be useful for pathogen identification. Our analyses led us to identify six main calorimetric parameters that can be used for the differentiation of species and identification of methicillin-resistant *S. aureus* isolates. These parameters are time to peak, time to activity, maximum metabolic rate, maximum metabolic velocity, and maximum metabolic and decay velocity (time). Time to peak was of particular interest to us, as it could potentially allow the diagnosis of these isolates in less than 7 h. Also, the fact that the time to peak is reduced in MRSA clinical isolates, underlines the possibility of a faster diagnosis of these strains in the clinics. However, these claims are based on a small sample size of MRSA strains and would need to be corroborated with a higher number of isolates.

Most of the calorimetric parameters derived from the thermograms relate to time, which is one of the most valuable variables in the nowadays-diagnostic scenarios. In addition, calorimetry has already proven to be effective in less than 24 h without sample manipulation, reducing considerably the current diagnostic time (5). By identifying universal parameters to distinguish resistance patterns in the microorganisms detected in a rapid manner, isothermal microcalorimetry could become one of the most powerful diagnostic tools for detecting and identifying infectious agents. Nowadays, the diagnostic tools are independent from the antimicrobial susceptibility testing techniques (23, 24), delaying the treatment of infectious diseases, where time plays the most important role to avoid the selection and spread of resistant microorganisms (3, 24).

We have also shown the metabolic differences between laboratory ATCC strains and clinical isolates by isothermal microcalorimetry (**Figure 4**). The similarities found between ATCC strains of both species suggest that the use of these strains for diagnostic research can lead to misidentification of strains. Laboratory strains are not representative of clinical isolates that come from patients with real infections and, therefore, simulations of these infections by using ATCC strains are not representative neither. This highlights the need of using the clinical isolates for the research and development of new diagnostic techniques. We propose here a workflow including the generation of model curves derived from the thermograms of clinical isolates, which would speed the detection of MRSA strains and, therefore improve the chances of their eradication (**Figure 5**). We have generated model curves using calorimetric data obtained from 84 clinical isolates of *S. aureus*, 7 of which were MRSA isolates, and 87 of *S. epidermidis*. These curves show a faster emergence of MRSA isolates compared to those from standard laboratory strains, highlighting the need of using clinical isolates for a more accurate identification process. Previous diagnostic studies for the molecular identification of MRSA and MSSA strains mostly base their work on reference strains (17, 18), however, there are other studies where clinical isolates started to be included (25-27). Nevertheless, isothermal microcalorimetry studies, as a more recent emerging tool, utilize primarily laboratory strains for the development of this technique (14-16), which underlines the need to include clinical isolates in future research for a more accurate diagnostic procedure.

Isothermal microcalorimetry seems to be a promising alternative to the current diagnostic tools used in the clinic; however, its early stage in development allows it just for now to complement the present techniques.

## CONTRIBUTIONS & ACKNOWLEDGEMENTS

FIAS and AT conceived the project. FIAS designed the experiments and protocol modifications. SZP performed the experiments. SZP and FIAS analyzed the experimental data. SZP and FIAS wrote the text. AT commented on the final manuscript.

## SUPPLEMENTARY INFORMATION

### Strain diversity Principal Component Analyses

We performed PCAs of the thermograms obtained from each strain. These analyses showed that a small portion of variance could be represented by each principal component generated (**Table S1**).

**Table S1.**
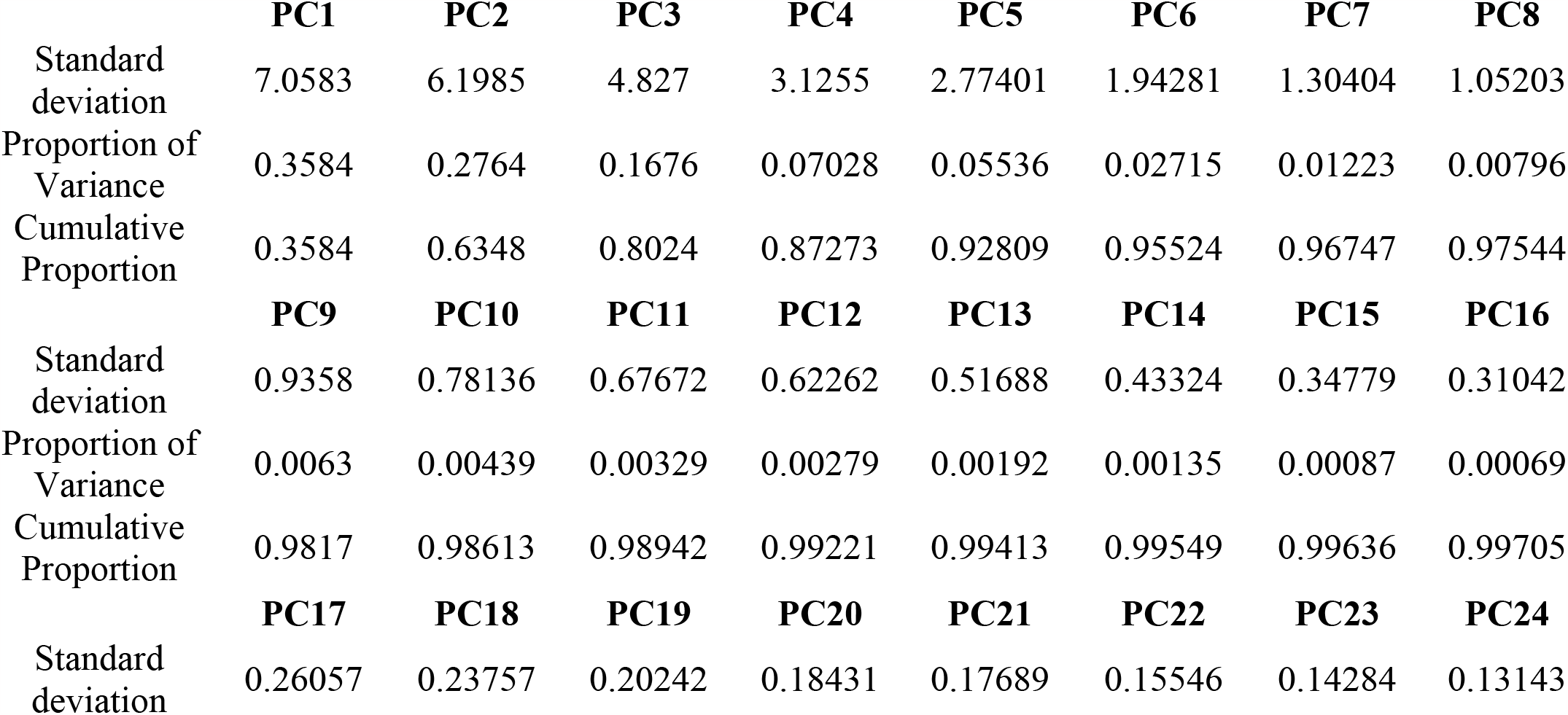
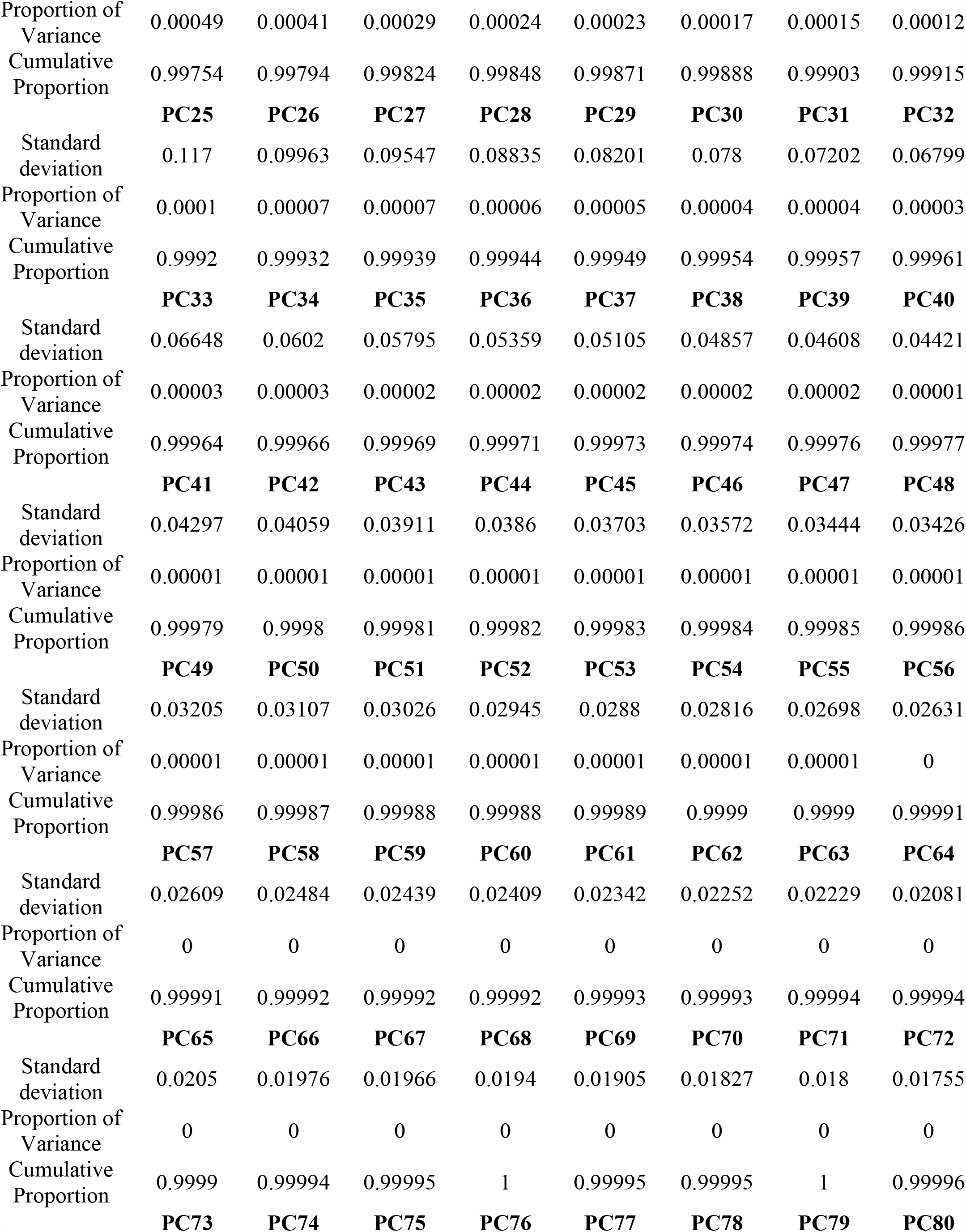
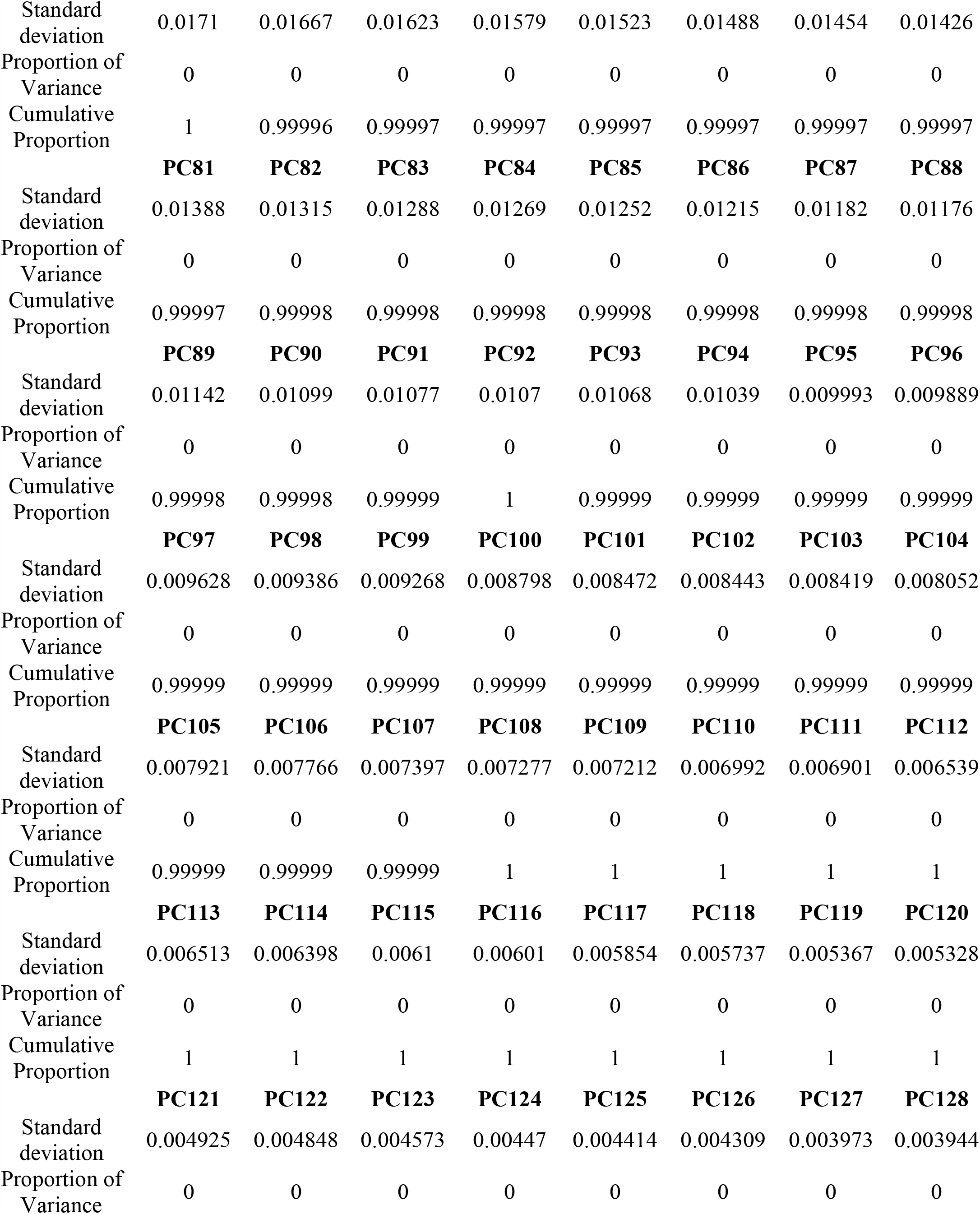
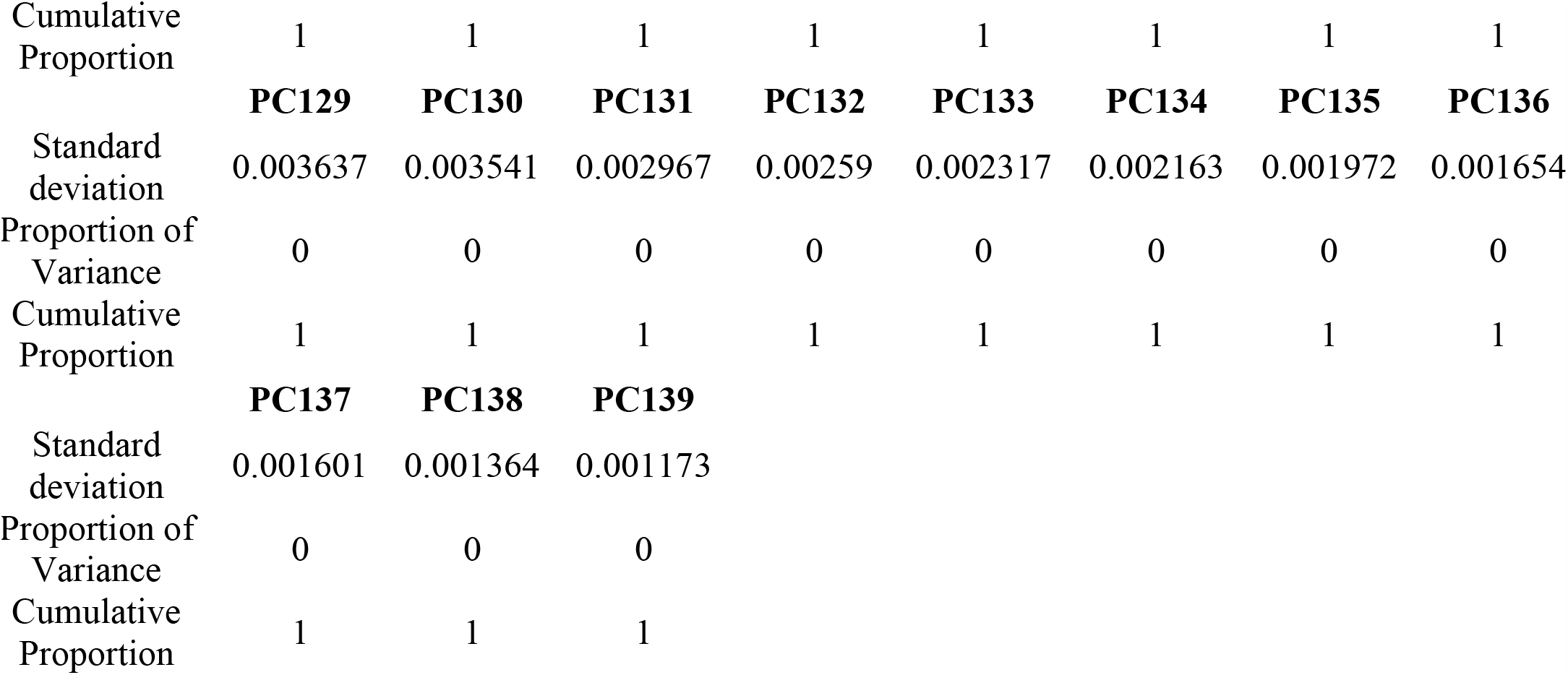
Importance of components. The cumulative portion of the first two components generated explain the 63.48% of the total variance.

### In-depth analysis of curve parameters: TSB thermograms have a higher resolution than MHB for both species

We used the online tool calData (Symcel) to collect the calorimetric parameters from each isolate. We performed MANOVA with all the parameters from each isolate that we obtained after growing them in TSB and MHB. We computed pairwise comparisons and parameters provided more relevant information about the calorimetric curves of bacteria when grown in TSB; more significant differences were identified in this medium than in MHB (**Table S2**). Therefore, further analyses were performed only with the data obtained from the TSB.

**Table S2.**
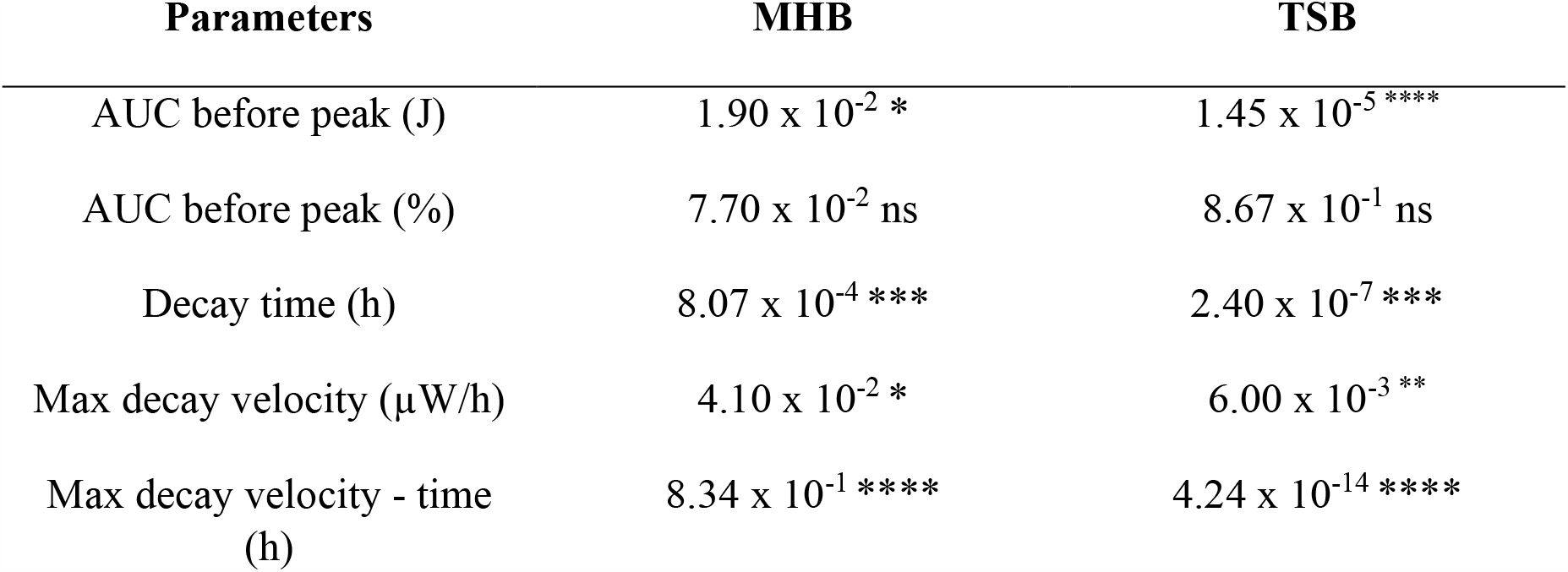
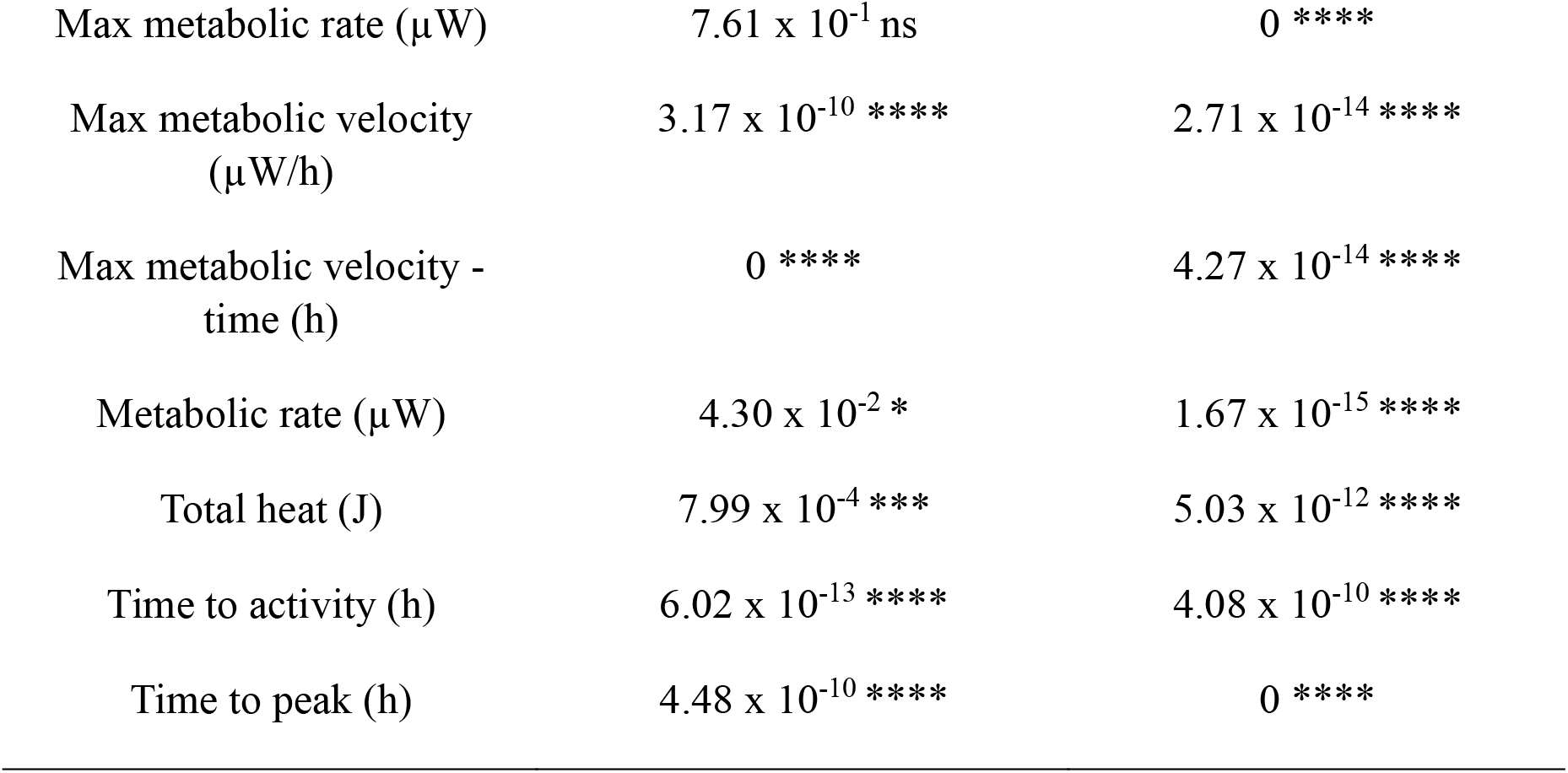
Adjusted p-values after the MANOVA with Bonferroni correction performed with all the calorimetric parameters obtained from each species in both media. More parameters (11/12) show significant differences (highlighted with asterisks) when tested in TSB than in MHB (10/12). Single asterisk denotes p < 0.05, double asterisks p < 0.01, triple asterisks p < 0.001, quadruple asterisks p < 0.0001, ns, not significant.

### Significant inter-species differences between relevant calorimetric parameters

We used Kruskal Wallis tests, followed by Wilcoxson signed-rank test with a Bonferroni correction and we performed pairwise comparisons of all the parameters of *S. aureus* and *S. epidermidis*. The majority of the comparisons were statistically significant (p<0.0001), except between decay time and time to activity, maximum decay velocity (time) and metabolic rate, and time to peak and metabolic rate (**Figure S1**).

**Figure S1.**
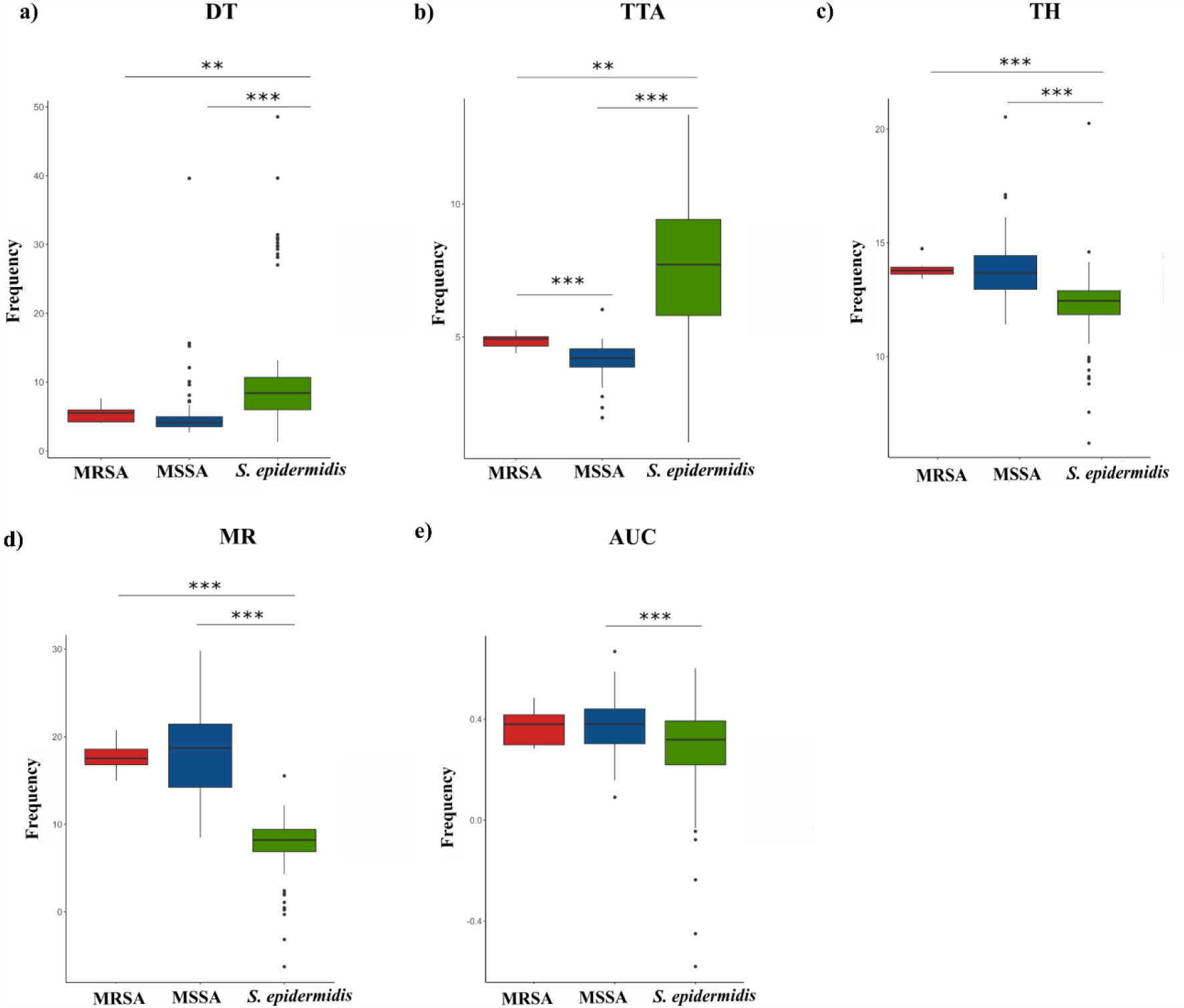
Pairwise comparisons of calorimetric parameters performed to study the differences between MRSA, MSSA and *S. epidermidis* isolates tested when grown in TSB. A total of 11 parameters were compared through Wilcoxson test with a Bonferroni adjustment. Significant differences between the parameters are represented by asterisks, where * represents p < 0.05, ** corresponds with a p < 0.01 and *** show a p < 0.001.

