## Supplementary material for "Towards faster identification of MRSA and MSSA: analysis of calorimetric curve parameters from large hospital bacterial collections"

### 1 Supplementary Tables and Figures

#### 1.1 Strain diversity Principal Component Analyses

We performed PCAs of the thermograms obtained from each strain. These analyses showed that a small portion of variance could be represented by each principal component generated (**Table S1**).

**Table S1. Importance of components.** The cumulative portion of the first two components generated explain the 63.48% of the total variance**.**

| | **PC1** | **PC2** | **PC3** | **PC4** | **PC5** | **PC6** | **PC7** | **PC8** |
|---|---|---|---|---|---|---|---|---|
| Standard deviation | 7.0583 | 6.1985 | 4.827 | 3.1255 | 2.77401 | 1.94281 | 1.30404 | 1.05203 |
| Proportion of Variance | 0.3584 | 0.2764 | 0.1676 | 0.07028 | 0.05536 | 0.02715 | 0.01223 | 0.00796 |
| Cumulative Proportion | 0.3584 | 0.6348 | 0.8024 | 0.87273 | 0.92809 | 0.95524 | 0.96747 | 0.97544 |
| | **PC9** | **PC10** | **PC11** | **PC12** | **PC13** | **PC14** | **PC15** | **PC16** |
| Standard deviation | 0.9358 | 0.78136 | 0.67672 | 0.62262 | 0.51688 | 0.43324 | 0.34779 | 0.31042 |
| Proportion of Variance | 0.0063 | 0.00439 | 0.00329 | 0.00279 | 0.00192 | 0.00135 | 0.00087 | 0.00069 |
| Cumulative Proportion | 0.9817 | 0.98613 | 0.98942 | 0.99221 | 0.99413 | 0.99549 | 0.99636 | 0.99705 |
| | **PC17** | **PC18** | **PC19** | **PC20** | **PC21** | **PC22** | **PC23** | **PC24** |
| Standard deviation | 0.26057 | 0.23757 | 0.20242 | 0.18431 | 0.17689 | 0.15546 | 0.14284 | 0.13143 |
| Proportion of Variance | 0.00049 | 0.00041 | 0.00029 | 0.00024 | 0.00023 | 0.00017 | 0.00015 | 0.00012 |
| Cumulative Proportion | 0.99754 | 0.99794 | 0.99824 | 0.99848 | 0.99871 | 0.99888 | 0.99903 | 0.99915 |
| | **PC25** | **PC26** | **PC27** | **PC28** | **PC29** | **PC30** | **PC31** | **PC32** |
| Standard deviation | 0.117 | 0.09963 | 0.09547 | 0.08835 | 0.08201 | 0.078 | 0.07202 | 0.06799 |
| Proportion of Variance | 0.0001 | 0.00007 | 0.00007 | 0.00006 | 0.00005 | 0.00004 | 0.00004 | 0.00003 |
| Cumulative Proportion | 0.9992 | 0.99932 | 0.99939 | 0.99944 | 0.99949 | 0.99954 | 0.99957 | 0.99961 |
| | **PC33** | **PC34** | **PC35** | **PC36** | **PC37** | **PC38** | **PC39** | **PC40** |
| Standard deviation | 0.06648 | 0.0602 | 0.05795 | 0.05359 | 0.05105 | 0.04857 | 0.04608 | 0.04421 |
| Proportion of Variance | 0.00003 | 0.00003 | 0.00002 | 0.00002 | 0.00002 | 0.00002 | 0.00002 | 0.00001 |
| Cumulative Proportion | 0.99964 | 0.99966 | 0.99969 | 0.99971 | 0.99973 | 0.99974 | 0.99976 | 0.99977 |

| | PC41 | PC42 | PC43 | PC44 | PC45 | PC46 | PC47 | PC48 |
|---|---|---|---|---|---|---|---|---|
| Standard deviation | 0.04297 | 0.04059 | 0.03911 | 0.0386 | 0.03703 | 0.03572 | 0.03444 | 0.03426 |
| Proportion of Variance | 0.00001 | 0.00001 | 0.00001 | 0.00001 | 0.00001 | 0.00001 | 0.00001 | 0.00001 |
| Cumulative Proportion | 0.99979 | 0.9998 | 0.99981 | 0.99982 | 0.99983 | 0.99984 | 0.99985 | 0.99986 |
| | **PC49** | **PC50** | **PC51** | **PC52** | **PC53** | **PC54** | **PC55** | **PC56** |
| Standard deviation | 0.03205 | 0.03107 | 0.03026 | 0.02945 | 0.0288 | 0.02816 | 0.02698 | 0.02631 |
| Proportion of Variance | 0.00001 | 0.00001 | 0.00001 | 0.00001 | 0.00001 | 0.00001 | 0.00001 | 0 |
| Cumulative Proportion | 0.99986 | 0.99987 | 0.99988 | 0.99988 | 0.99989 | 0.9999 | 0.9999 | 0.99991 |
| | **PC57** | **PC58** | **PC59** | **PC60** | **PC61** | **PC62** | **PC63** | **PC64** |
| Standard deviation | 0.02609 | 0.02484 | 0.02439 | 0.02409 | 0.02342 | 0.02252 | 0.02229 | 0.02081 |
| Proportion of Variance | 0 | 0 | 0 | 0 | 0 | 0 | 0 | 0 |
| Cumulative Proportion | 0.99991 | 0.99992 | 0.99992 | 0.99992 | 0.99993 | 0.99993 | 0.99994 | 0.99994 |
| | **PC65** | **PC66** | **PC67** | **PC68** | **PC69** | **PC70** | **PC71** | **PC72** |
| Standard deviation | 0.0205 | 0.01976 | 0.01966 | 0.0194 | 0.01905 | 0.01827 | 0.018 | 0.01755 |
| Proportion of Variance | 0 | 0 | 0 | 0 | 0 | 0 | 0 | 0 |
| Cumulative Proportion | 0.9999 | 0.99994 | 0.99995 | 1 | 0.99995 | 0.99995 | 1 | 0.99996 |
| | **PC73** | **PC74** | **PC75** | **PC76** | **PC77** | **PC78** | **PC79** | **PC80** |
| Standard deviation | 0.0171 | 0.01667 | 0.01623 | 0.01579 | 0.01523 | 0.01488 | 0.01454 | 0.01426 |
| Proportion of Variance | 0 | 0 | 0 | 0 | 0 | 0 | 0 | 0 |
| Cumulative Proportion | 1 | 0.99996 | 0.99997 | 0.99997 | 0.99997 | 0.99997 | 0.99997 | 0.99997 |
| | **PC81** | **PC82** | **PC83** | **PC84** | **PC85** | **PC86** | **PC87** | **PC88** |
| Standard deviation | 0.01388 | 0.01315 | 0.01288 | 0.01269 | 0.01252 | 0.01215 | 0.01182 | 0.01176 |
| Proportion of Variance | 0 | 0 | 0 | 0 | 0 | 0 | 0 | 0 |
| Cumulative Proportion | 0.99997 | 0.99998 | 0.99998 | 0.99998 | 0.99998 | 0.99998 | 0.99998 | 0.99998 |
| | **PC89** | **PC90** | **PC91** | **PC92** | **PC93** | **PC94** | **PC95** | **PC96** |
| Standard deviation | 0.01142 | 0.01099 | 0.01077 | 0.0107 | 0.01068 | 0.01039 | 0.009993 | 0.009889 |

| | | | | | | | | |
|---|---|---|---|---|---|---|---|---|
| Proportion of Variance | 0 | 0 | 0 | 0 | 0 | 0 | 0 | 0 |
| Cumulative Proportion | 0.99998 | 0.99998 | 0.99999 | 1 | 0.99999 | 0.99999 | 0.99999 | 0.99999 |
| | **PC97** | **PC98** | **PC99** | **PC100** | **PC101** | **PC102** | **PC103** | **PC104** |
| Standard deviation | 0.009628 | 0.009386 | 0.009268 | 0.008798 | 0.008472 | 0.008443 | 0.008419 | 0.008052 |
| Proportion of Variance | 0 | 0 | 0 | 0 | 0 | 0 | 0 | 0 |
| Cumulative Proportion | 0.99999 | 0.99999 | 0.99999 | 0.99999 | 0.99999 | 0.99999 | 0.99999 | 0.99999 |
| | **PC105** | **PC106** | **PC107** | **PC108** | **PC109** | **PC110** | **PC111** | **PC112** |
| Standard deviation | 0.007921 | 0.007766 | 0.007397 | 0.007277 | 0.007212 | 0.006992 | 0.006901 | 0.006539 |
| Proportion of Variance | 0 | 0 | 0 | 0 | 0 | 0 | 0 | 0 |
| Cumulative Proportion | 0.99999 | 0.99999 | 0.99999 | 1 | 1 | 1 | 1 | 1 |
| | **PC113** | **PC114** | **PC115** | **PC116** | **PC117** | **PC118** | **PC119** | **PC120** |
| Standard deviation | 0.006513 | 0.006398 | 0.0061 | 0.00601 | 0.005854 | 0.005737 | 0.005367 | 0.005328 |
| Proportion of Variance | 0 | 0 | 0 | 0 | 0 | 0 | 0 | 0 |
| Cumulative Proportion | 1 | 1 | 1 | 1 | 1 | 1 | 1 | 1 |
| | **PC121** | **PC122** | **PC123** | **PC124** | **PC125** | **PC126** | **PC127** | **PC128** |
| Standard deviation | 0.004925 | 0.004848 | 0.004573 | 0.00447 | 0.004414 | 0.004309 | 0.003973 | 0.003944 |
| Proportion of Variance | 0 | 0 | 0 | 0 | 0 | 0 | 0 | 0 |
| Cumulative Proportion | 1 | 1 | 1 | 1 | 1 | 1 | 1 | 1 |
| | **PC129** | **PC130** | **PC131** | **PC132** | **PC133** | **PC134** | **PC135** | **PC136** |
| Standard deviation | 0.003637 | 0.003541 | 0.002967 | 0.00259 | 0.002317 | 0.002163 | 0.001972 | 0.001654 |
| Proportion of Variance | 0 | 0 | 0 | 0 | 0 | 0 | 0 | 0 |
| Cumulative Proportion | 1 | 1 | 1 | 1 | 1 | 1 | 1 | 1 |
| | **PC137** | **PC138** | **PC139** | | | | | |
| Standard deviation | 0.001601 | 0.001364 | 0.001173 | | | | | |
| Proportion of Variance | 0 | 0 | 0 | | | | | |
| Cumulative Proportion | 1 | 1 | 1 | | | | | |

### 1.2 In-depth analysis of curve parameters: TSB thermograms have a higher resolution than MHB for both species

We used the online tool calData (Symcel) to collect the calorimetric parameters from each isolate. We performed MANOVA with all the parameters from each isolate that we obtained after growing them in TSB and MHB. We computed pairwise comparisons and parameters provided more relevant information about the calorimetric curves of bacteria when grown in TSB; more significant differences were identified in this medium than in MHB (**Table S2**). Therefore, further analyses were performed only with the data obtained from the TSB.

**Table S2. Adjusted p-values after the MANOVA with Bonferroni correction performed with all the calorimetric parameters obtained from each species in both media.** More parameters (11/12) show significant differences (highlighted with asterisks) when tested in TSB than in MHB (10/12). Single asterisk denotes $p < 0.05$, double asterisks $p < 0.01$, triple asterisks $p < 0.001$, quadruple asterisks $p < 0.0001$, ns, not significant.

| Parameters | MHB | TSB |
|---|---|---|
| AUC before peak (J) | $1.90 \times 10^{-2}$ * | $1.45 \times 10^{-5}$ **** |
| AUC before peak (%) | $7.70 \times 10^{-2}$ ns | $8.67 \times 10^{-1}$ ns |
| Decay time (h) | $8.07 \times 10^{-4}$ *** | $2.40 \times 10^{-7}$ *** |
| Max decay velocity (µW/h) | $4.10 \times 10^{-2}$ * | $6.00 \times 10^{-3}$ ** |
| Max decay velocity - time (h) | $8.34 \times 10^{-1}$ **** | $4.24 \times 10^{-14}$ **** |
| Max metabolic rate (µW) | $7.61 \times 10^{-1}$ ns | 0 **** |
| Max metabolic velocity (µW/h) | $3.17 \times 10^{-10}$ **** | $2.71 \times 10^{-14}$ **** |
| Max metabolic velocity - time (h) | 0 **** | $4.27 \times 10^{-14}$ **** |
| Metabolic rate (µW) | $4.30 \times 10^{-2}$ * | $1.67 \times 10^{-15}$ **** |
| Total heat (J) | $7.99 \times 10^{-4}$ *** | $5.03 \times 10^{-12}$ **** |
| Time to activity (h) | $6.02 \times 10^{-13}$ **** | $4.08 \times 10^{-10}$ **** |
| Time to peak (h) | $4.48 \times 10^{-10}$ **** | 0 **** |

### 1.3 Significant inter-species differences between relevant calorimetric parameters

We used Kruskal Wallis tests, followed by Wilcoxson signed-rank test with a Bonferroni correction and we performed pairwise comparisons of all the parameters of *S. aureus* and *S. epidermidis*. The majority of the comparisons were statistically significant (p<0.0001), except between decay time and time to activity, maximum decay velocity (time) and metabolic rate, and time to peak and metabolic rate (**Figure S1**).

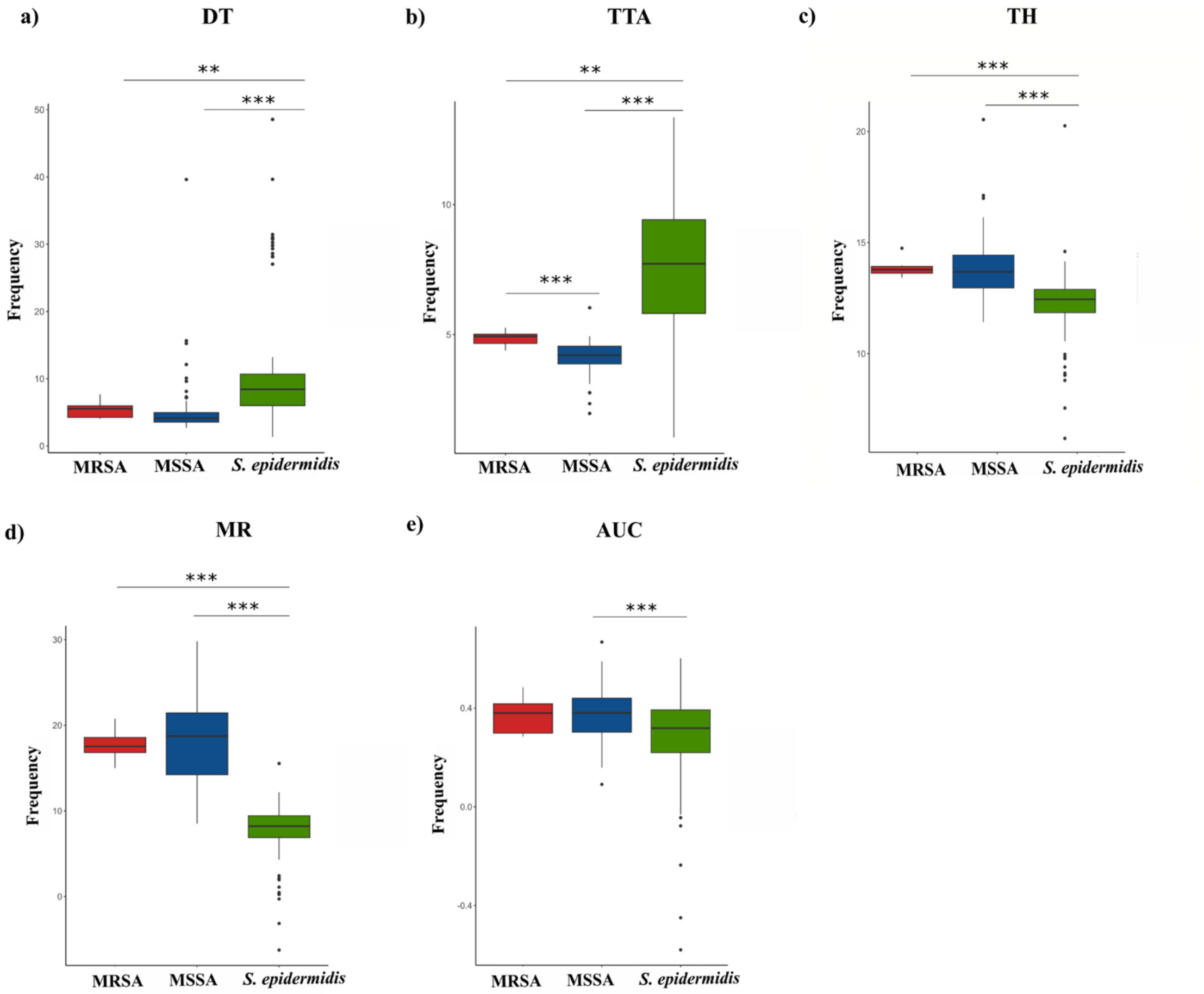


**Figure S1. Pairwise comparisons of calorimetric parameters performed to study the differences between MRSA, MSSA and *S. epidermidis* isolates tested when grown in TSB.** A total of 11 parameters were compared through Wilcoxson test with a Bonferroni adjustment. Significant differences between the parameters are represented by asterisks, where * represents $p < 0.05$, ** corresponds with a $p < 0.01$ and *** show a $p < 0.001$.
